## Supplemental Information for "How hibernation in frogs drives brain and reproductive evolution in opposite directions"

This file includes:

Figures S1 to S10
Tables S1 to S17

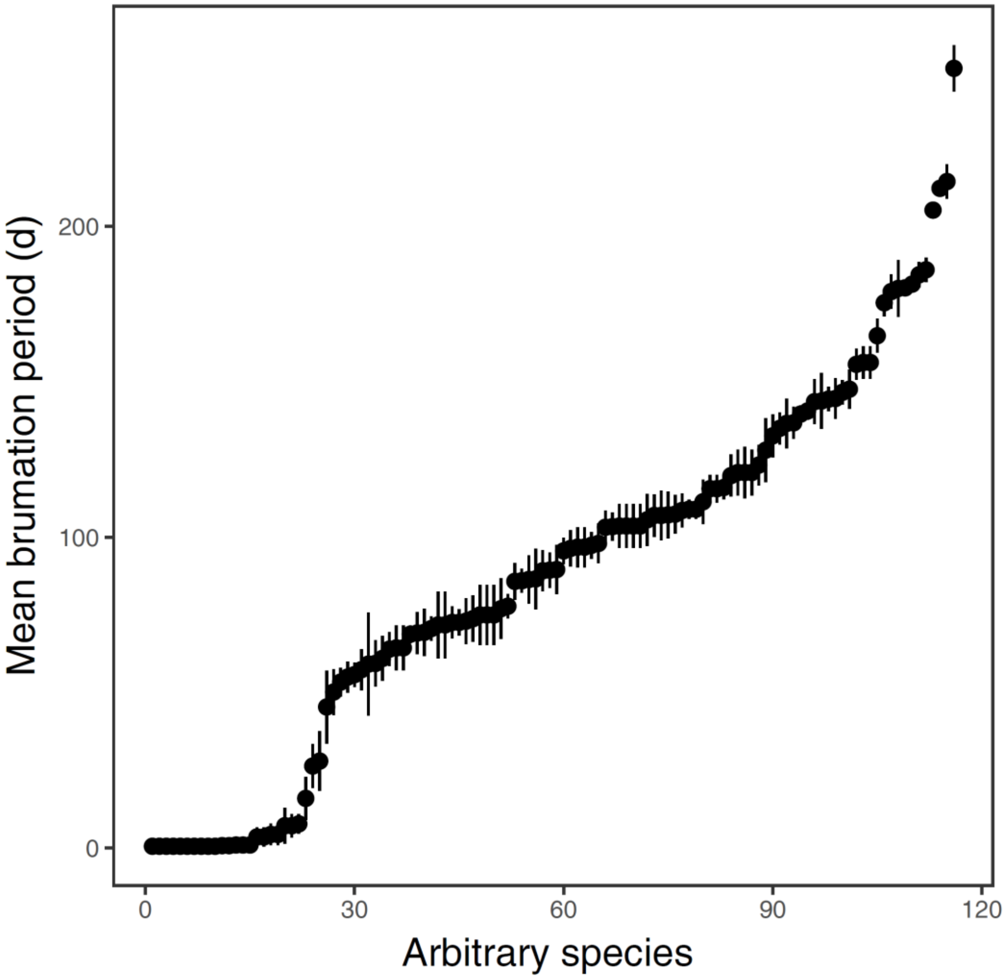

**Fig. S1.** Distribution of brumation durations (in days) across the 116 anuran species. Each dot represents the species-specific mean ± 1 SE of the 5 years (2012−2016) that specimens were collected. For each year and species, all days with a collection site-specific ambient temperature below the experimentally quantified, species-specific temperature thresholds were summed to estimate the brumation period.

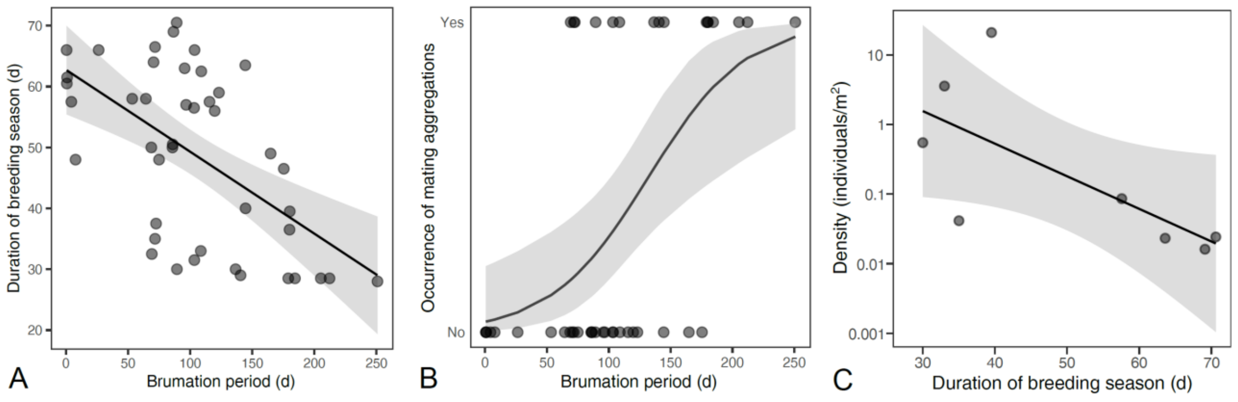

**Fig. S2.** Effects of brumation on breeding conditions. Prolonged brumation shortened the breeding season (**A**), thereby increasing the probability of dense breeding aggregations (**B**). This effect was also supported by a trend toward higher mean population densities in species with a shorter breeding season (**C**).

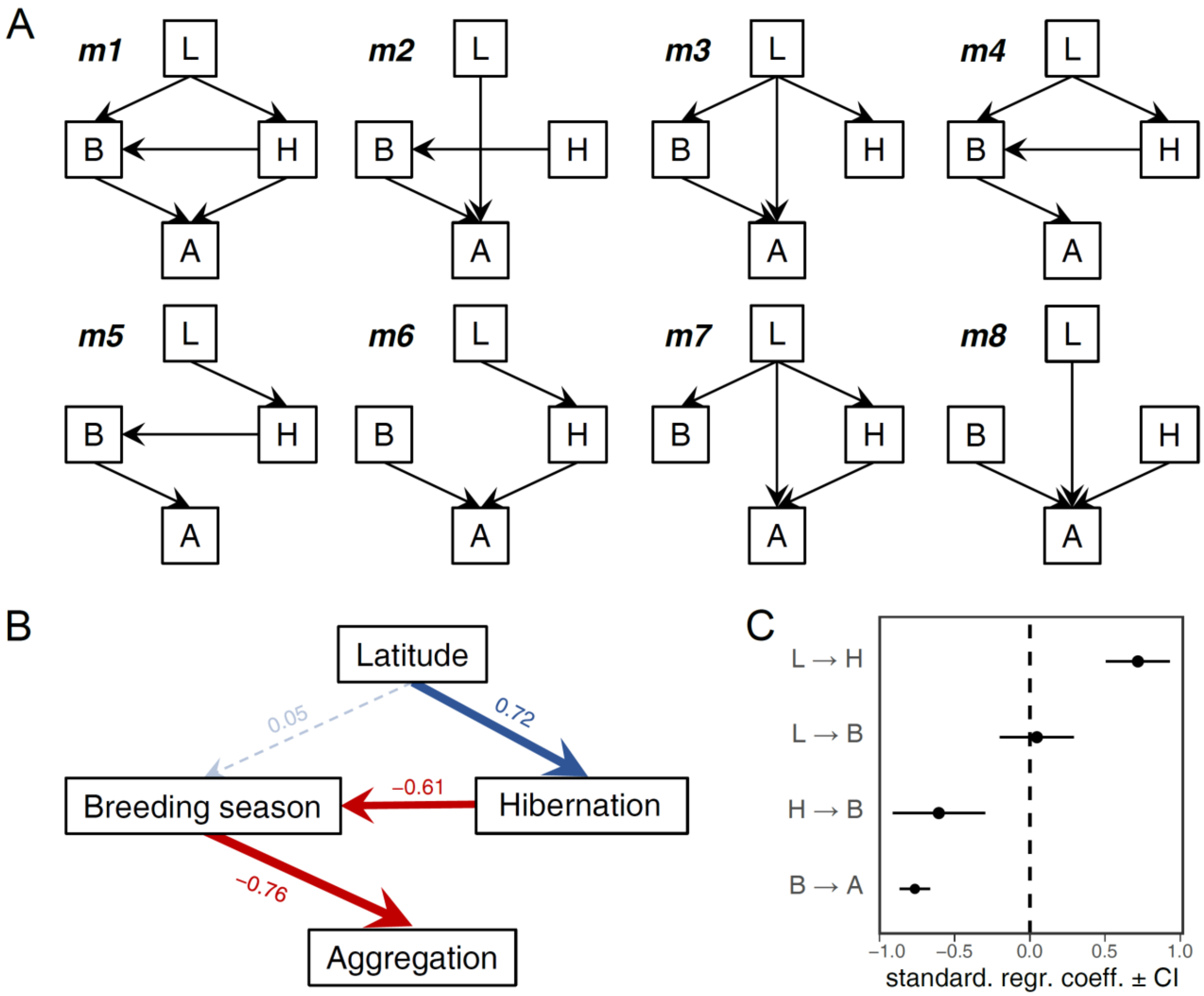

**Fig. S3.** Confirmatory path analysis on the links between environmental variation, the brumation duration and breeding activity and the formation of breeding aggregations. Directed acyclic graphs representing 8 candidate models (**A**) formed the predictions for the path analysis, resulting in the average path model (**B**). The path coefficients (standardised regression coefficients) and their 95% confidence intervals are depicted in **C**. The variable letters correspond to the first letters of the full variable names in panel **B**. The model comparisons are listed in Table S13. The results were qualitatively identical when substituting latitude with elevation, mean temperature, or the coefficient of variation in temperature (except that the effect of mean temperature on brumation duration was negative as predicted). Note that here we used ‘hibernation’ instead of ‘brumation’ to avoid conflicts in codes.

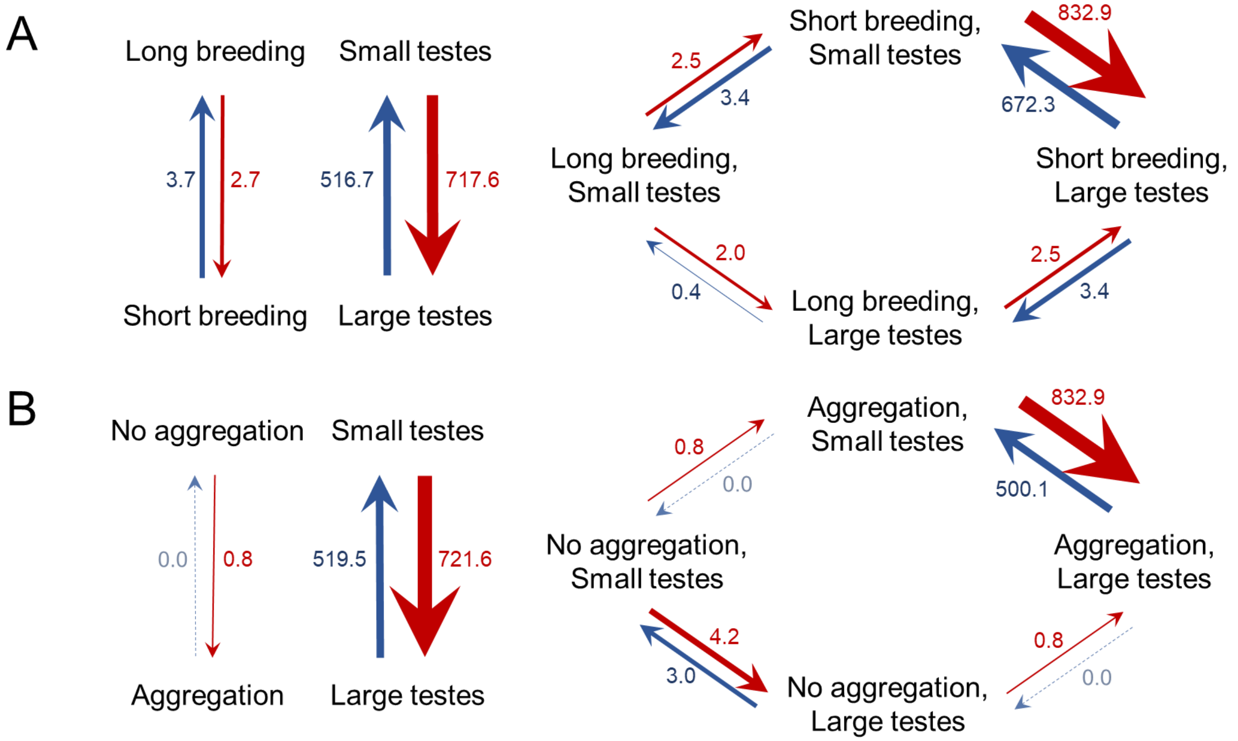

**Fig. S4.** Directional test of trait evolution between relative testis size and breeding conditions. Transition rates between binary states for relative testis size and (**A**) the duration of the breeding season or (**B**) aggregation formation. Independent models are shown on the left, models with changes in relative testis size dependent on the breeding parameters are depicted on the right. For both pairs of variables, the independent model had the highest support (i.e., lowest AIC scores and highest AIC weights: **A**: $w_{AIC} = 0.50$, B: $w_{AIC} = 0.66$), but the dependent models shown here were either not significantly less supported (**A**: $\Delta AIC = 0.83$, $w_{AIC} = 0.33$) or relatively weakly so (**B**: $\Delta AIC = 2.38$, $w_{AIC} = 0.20$). The remaining two models (changes in breeding parameters depending on those in relative testis size or both variables evolving interdependently) found much less support in both analyses (**A**: $\Delta AIC \geq 3.40$, $w_{AIC} \leq 0.09$, **B**: $\Delta AIC \geq 3.64$, $w_{AIC} \leq 0.11$).

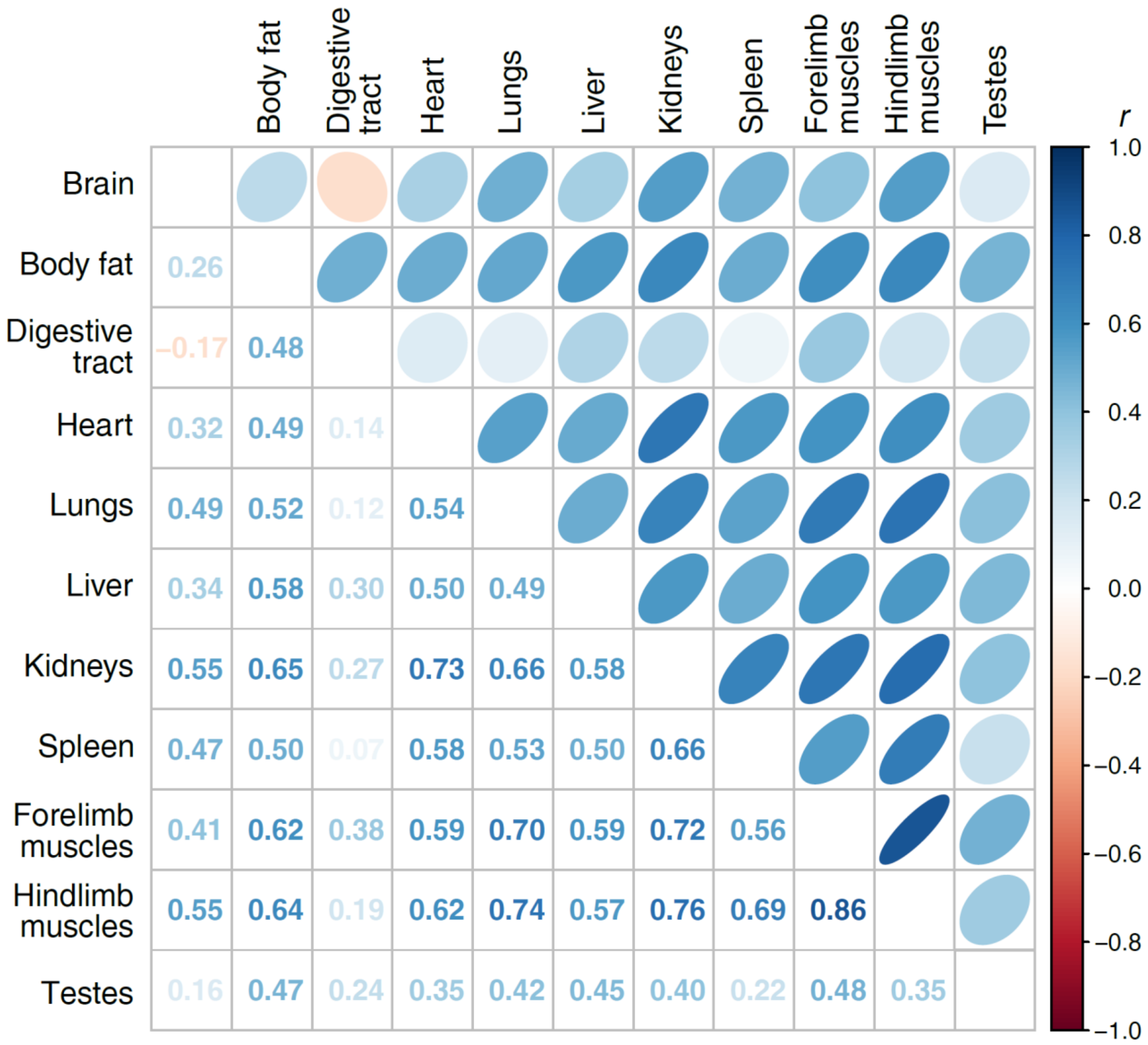

**Fig. S5.** Pairwise partial correlations between the different tissues. All correlations are controlled for snout-vent length and phylogeny, expressed both by ellipses (above diagonal) and the correlation coefficients (below the diagonal). All correlations with $|r_p| \geq 0.19$ had $P < 0.05$ before and after accounting for multiple testing based on the False Discovery Rate.

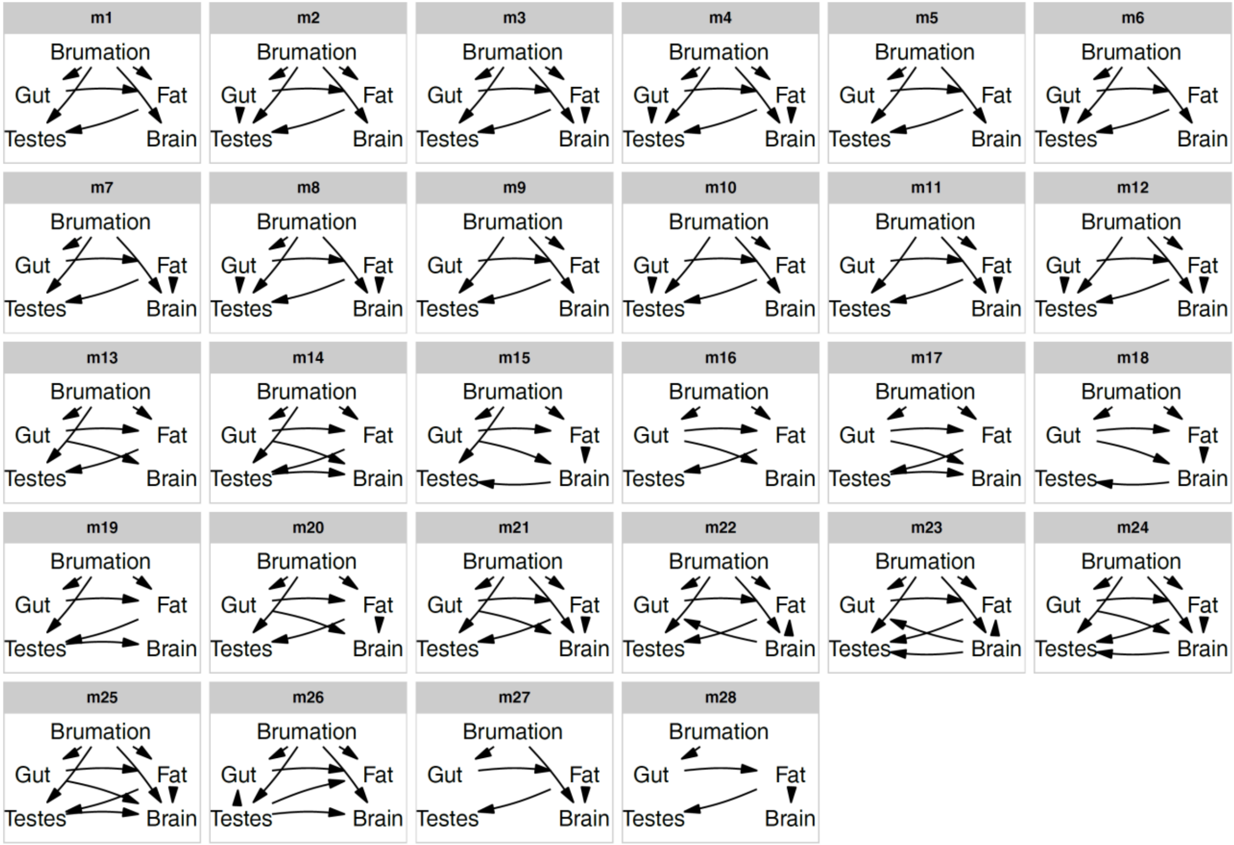

**Fig. S6.** Candidate path models. Directed acyclic graphs representing 28 candidate models that were compared to disentangle the relationships between five traits through a phylogenetic confirmatory path analysis and multi-model inference. For spatial reasons, the digestive tract is termed gut, and we omitted the paths from snout-vent length to all five variables in each panel to control tissue masses for body size or to improve *d*-separation (in the case of brumation).

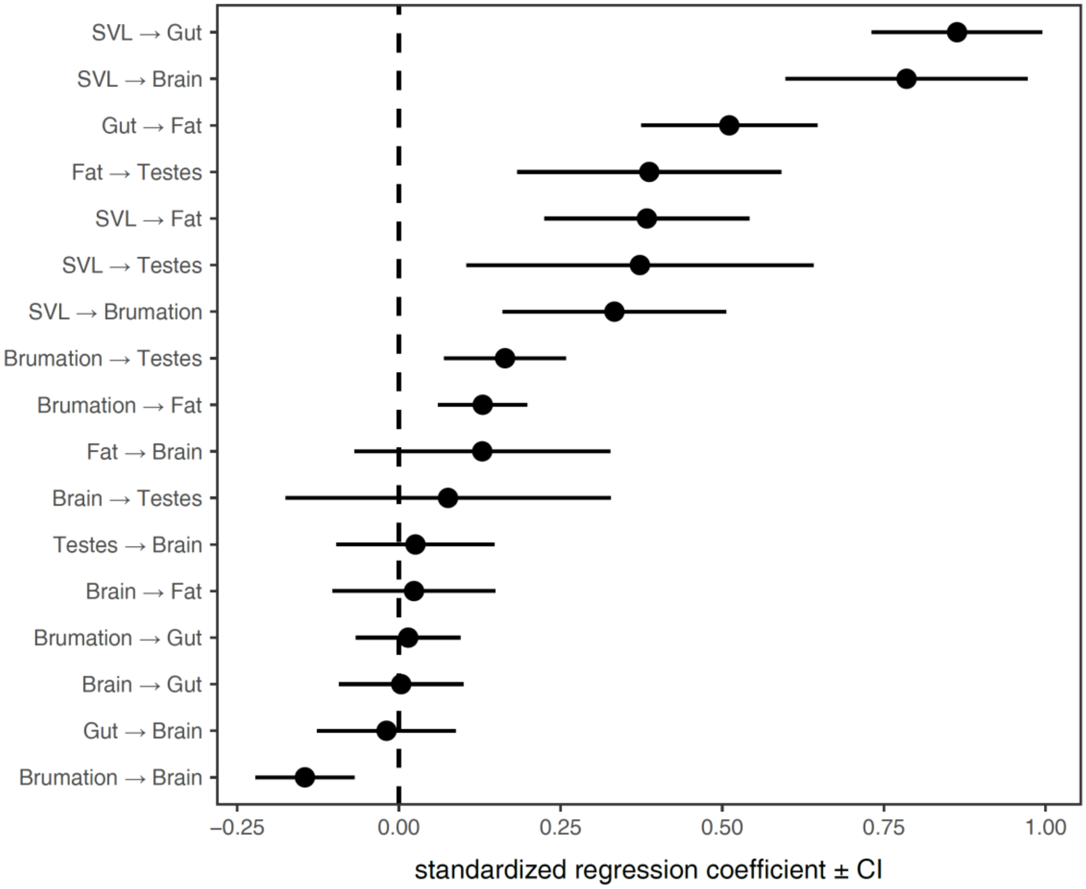

**Fig. S7.** Coefficients of the averaged phylogenetic path model. The path coefficients (standardised regression coefficients) are depicted with their 95% confidence intervals, corresponding to the path analysis in Fig. 4 (directed path models in Fig. S6). The paths from SVL to other traits are included only to control for body size (in the case of tissues) or to improve *d*-separation (SVL → Brumation). For visual clarity, these are omitted in the directed acyclic graphs (Fig. S6) and the final averaged model (Fig. 4).

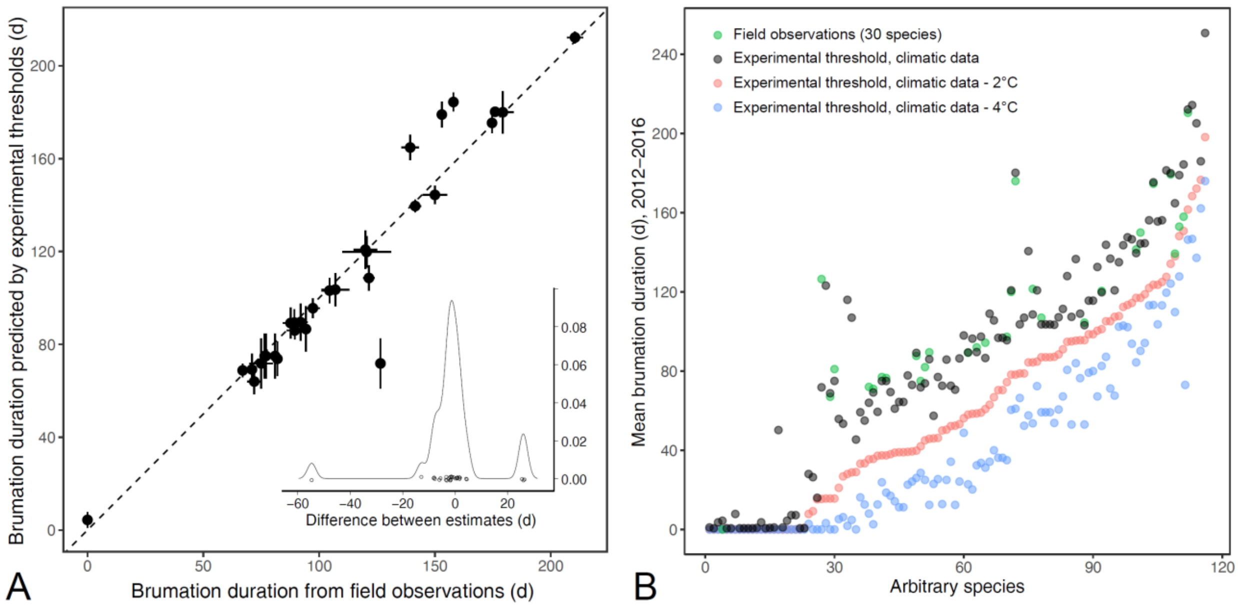

**Fig. S8.** Validation of the different estimates of brumation duration. **A:** Comparisons of field observations with predicted brumation periods based on experimental thermal thresholds and site-specific climate data. The dashed line represents unity. The inserted density plot indicates that twenty-five of the 30 estimated values were within ≤8 days of the field observations across the spectrum, with the remaining five estimates deviating by approx. 13, 26 (3×) or, for one outlier, 55 days. **B:** Variation in species-specific brumation periods based on different approaches. The green dots are the same field-based data as in panel **A**. The black dots represent values that combine the experimental temperature thresholds with the site-specific climatic data (predicted brumation below the threshold). The red and blue dots are based on the same experimental thresholds, but this time following the air temperature profiles of 2 and 4°C below these thresholds, respectively, to account for the potential buffering effect of the soil for underground hibernacula. Since the green dots were most closely associated with the black ones (i.e., without the additional buffer), we focused primarily on those values, but conducted additional analyses using the shorter, buffered brumation periods with no qualitative change. Finally, the horizontal dashed line splits the species into those with brumation likely present (above) or absent (below) to generate binary brumation variables for more direct comparison with previous studies in other taxa. This cut-off of ≤40 *versus* >40 days was motivated by the relatively large gap in the sequence of brumation periods in both the black and blue dots and that anurans are likely to survive relatively short cold spells without special adaptations. This cut-off generated divergent datasets between the direct experimental thresholds (black: $N$ = 91 species with and 25 without prolonged brumation) and buffered thresholds (blue: $N$ = 47 species with and 69 without prolonged brumation).

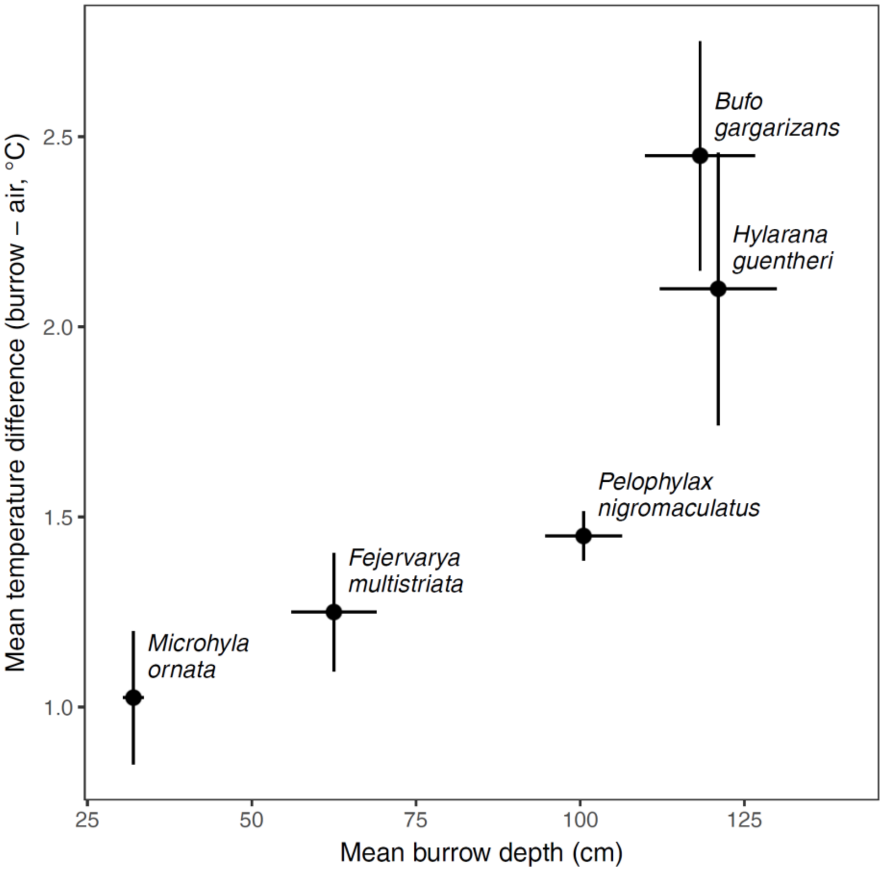

**Fig. S9.** Comparison of air and burrow temperatures. The difference between burrow and outside air temperature increased with burrow depth across the five species examined in a pilot study. The points reflect the means with standard errors around them based on four burrows per species.

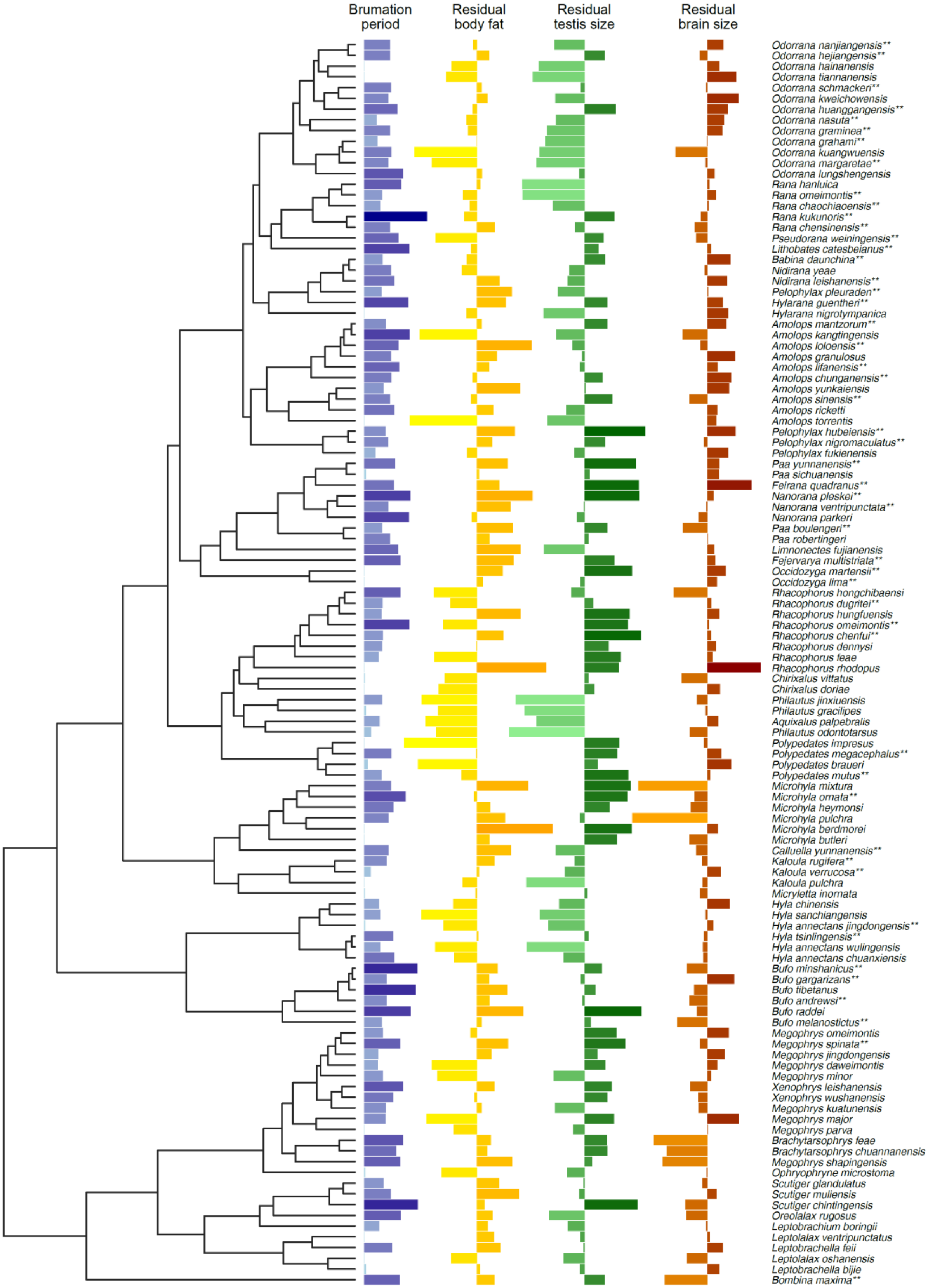

**Fig. S10.** Phylogeny of the species used in the study. Species with data on pre-brumation condition are labelled with double-asterisks (**). Residual trait sizes are derived from log-log linear regressions against snout-vent length.

**Table S1.** Within-species repeatability in male morphology, climatic variables, and brumation duration. CV = coefficient of variation, P2T = duration of the dry season (in months), with a month defined as dry when its total precipitation is less than two times the mean temperature. All analyses are based on the males (in breeding condition) or specific sampling locations for *N* = 116 species, except field observations of brumation duration (*N* = 30 species).

| Variable | *N* | *R [95%CI]* | *P* |
|---|---|---|---|
| *Male tissue masses* | | | |
| SVL | 396 | 0.950 [0.933, 0.962] | <0.001 |
| Brain | 396 | 0.948 [0.930, 0.961] | <0.001 |
| Body fat | 396 | 0.958 [0.942, 0.969] | <0.001 |
| Digestive tract | 396 | 0.949 [0.931, 0.962] | <0.001 |
| Heart | 396 | 0.918 [0.890, 0.939] | <0.001 |
| Lungs | 396 | 0.920 [0.890, 0.939] | <0.001 |
| Liver | 396 | 0.915 [0.885, 0.936] | <0.001 |
| Kidneys | 396 | 0.921 [0.893, 0.941] | <0.001 |
| Spleen | 396 | 0.937 [0.914, 0.952] | <0.001 |
| Forelimb muscles | 396 | 0.947 [0.929, 0.960] | <0.001 |
| Hindlimb muscles | 396 | 0.961 [0.946, 0.971] | <0.001 |
| Testes | 396 | 0.947 [0.929, 0.961] | <0.001 |
| *Climate variables* | | | |
| Mean temperature | 580 | 0.982 [0.976, 0.986] | <0.001 |
| Mean precipitation | 580 | 0.757 [0.693, 0.802] | <0.001 |
| CV temperature | 580 | 0.978 [0.971, 0.982] | <0.001 |
| CV precipitation | 580 | 0.401 [0.327, 0.486] | <0.001 |
| P2T | 580 | 0.688 [0.614, 0.751] | <0.001 |
| *Brumation* | | | |
| Brumation duration (estimated from lab data) | 580 | 0.947 [0.928, 0.959] | <0.001 |
| Brumation duration (field observations) | 66 | 0.982 [0.965, 0.991] | <0.001 |
| *Breeding season* | | | |
| Breeding season duration | 86 | 0.881 [0.785, 0.936] | <0.001 |

**Table S2.** Species-specific environmental effects on the brumation duration. CV = coefficient of variation, P2T = duration of the dry season (in months), with a month defined as dry when its total precipitation is less than two times the mean temperature. All analyses are based on df = 114, and confidence intervals are bootstrapped (*N* = 100 simulations). λ = phylogenetic scaling parameter.

| Predictor | *β* [95%CI] | *r* [95%CI] | *t* | *P* | *λ* [95%CI] |
|---|---|---|---|---|---|
| Latitude | 0.72 [ 0.61, 0.82] | 0.72 [ 0.63, 0.78] | 11.03 | **<0.001** | 0.00 [0.00, 0.10] |
| Longitude | 0.09 [-0.06, 0.28] | 0.09 [-0.09, 0.27] | 1.00 | 0.321 | 0.47 [0.00, 0.69] |
| Elevation | 0.43 [ 0.24, 0.62] | 0.43 [ 0.28, 0.56] | 5.15 | **<0.001** | 0.00 [0.00, 0.10] |
| Mean temperature | -0.56 [-0.69, -0.38] | -0.56 [-0.66, -0.42] | -7.14 | **<0.001** | 0.00 [0.00, 0.11] |
| Mean precipitation | -0.30 [-0.45, -0.13] | -0.30 [-0.45, -0.13] | -3.42 | **<0.001** | 0.00 [0.00, 0.10] |
| CV temperature | 0.65 [ 0.51, 0.78] | 0.65 [ 0.54, 0.73] | 9.08 | **<0.001** | 0.00 [0.00, 0.11] |
| CV precipitation | 0.01 [-0.18, 0.21] | 0.01 [-0.17, 0.19] | 0.06 | 0.948 | 0.42 [0.00, 0.65] |
| P2T | -0.37 [-0.52, -0.19] | -0.38 [-0.51, -0.21] | -4.38 | **<0.001** | 0.42 [0.00, 0.66] |

**Table S3.** Species-specific environmental effects on the brumation duration, excluding those 25 species that are unlikely to hibernate (≤27 days). CV = coefficient of variation, P2T = duration of the dry season (in months), with a month defined as dry when its total precipitation is less than two times the mean temperature. All analyses are based on df = 89, and confidence intervals are bootstrapped (*N* = 100 simulations). λ = phylogenetic scaling parameter.

| Predictor | *β* [95%CI] | *r* [95%CI] | *t* | *P* | *λ* [95%CI] |
|---|---|---|---|---|---|
| Latitude | 0.46 [ 0.28, 0.65] | 0.46 [ 0.29, 0.60] | 4.94 | **<0.001** | 0.00 [0.00, 0.11] |
| Longitude | -0.15 [-0.35, 0.08] | -0.15 [-0.33, 0.06] | -1.38 | 0.170 | 0.00 [0.00, 0.07] |
| Elevation | 0.42 [ 0.21, 0.61] | 0.42 [ 0.24, 0.56] | 4.38 | **<0.001** | 0.00 [0.00, 0.11] |
| Mean temperature | -0.36 [-0.58, -0.15] | -0.36 [-0.52, -0.17] | -3.69 | **<0.001** | 0.00 [0.00, 0.10] |
| Mean precipitation | -0.35 [-0.51, -0.12] | -0.35 [-0.51, -0.16] | -3.54 | **<0.001** | 0.00 [0.00, 0.08] |
| CV temperature | 1.17 [ 0.47, 1.80] | 0.72 [ 0.28, 0.87] | 3.42 | **0.006** | 0.79 [0.00, 1.00] |
| CV precipitation | 0.03 [-0.16, 0.25] | 0.03 [-0.18, 0.23] | 0.27 | 0.785 | 0.00 [0.00, 0.06] |
| P2T | -0.21 [-0.38, -0.01] | -0.21 [-0.39, 0.00] | -2.02 | **0.046** | 0.00 [0.00, 0.09] |

**Table S4.** Species-specific environmental effects on the temperature at which males entered or left their inactive state. All analyses are limited to those 91 species that are likely to overwinter (≥47 days below experimental temperature threshold), with df = 89, and confidence intervals are bootstrapped (*N* = 100 simulations). CV = coefficient of variation, P2T = duration of the dry season (in months), with a month defined as dry when its total precipitation is less than two times the mean temperature, λ = phylogenetic scaling parameter.

| Predictor | *β* [95%CI] | *r* [95%CI] | *t* | *P* | *λ* |
|---|---|---|---|---|---|
| *Entering inactive state:* | | | | | |
| Latitude | -0.10 [-0.26, 0.09] | -0.12 [-0.31, 0.09] | -1.16 | 0.247 | 0.32 [0.00, 0.56] |
| Longitude | 0.24 [ 0.09, 0.41] | 0.31 [ 0.11, 0.47] | 3.02 | **0.003** | 0.00 [0.00, 0.07] |
| Elevation | -1.42 [-1.94, -0.99] | -0.51 [-0.63, -0.34] | -5.56 | **<0.001** | 0.07 [0.00, 0.23] |
| Mean temperature | 0.24 [ 0.13, 0.35] | 0.46 [ 0.28, 0.59] | 4.84 | **<0.001** | 0.00 [0.00, 0.10] |
| Mean precipitation | 0.01 [-0.01, 0.03] | 0.07 [-0.13, 0.27] | 0.71 | 0.480 | 0.39 [0.00, 0.61] |
| CV temperature | -1.78 [-2.69, -0.88] | -0.39 [-0.54, -0.20] | -4.03 | **<0.001** | 0.00 [0.00, 0.07] |
| CV precipitation | -3.62 [-6.87, 0.10] | -0.21 [-0.39, 0.00] | -1.99 | **0.049** | 0.00 [0.00, 0.06] |
| P2T | 0.35 [ 0.01, 0.63] | 0.22 [ 0.02, 0.40] | 2.18 | **0.032** | 0.43 [0.00, 0.66] |
| *Leaving inactive state:* | | | | | |
| Latitude | -0.12 [-0.30, 0.08] | -0.14 [-0.33, 0.07] | -1.34 | 0.184 | 0.47 [0.00, 0.68] |
| Longitude | 0.28 [ 0.12, 0.46] | 0.35 [ 0.15, 0.50] | 3.48 | **<0.001** | 0.37 [0.00, 0.59] |
| Elevation | -1.63 [-2.13, -1.18] | -0.55 [-0.66, -0.39] | -6.23 | **<0.001** | 0.23 [0.00, 0.46] |
| Mean temperature | 0.21 [ 0.09, 0.32] | 0.39 [ 0.20, 0.54] | 3.97 | **<0.001** | 0.35 [0.00, 0.59] |
| Mean precipitation | 0.01 [ 0.00, 0.03] | 0.14 [-0.07, 0.33] | 1.32 | 0.191 | 0.47 [0.00, 0.68] |
| CV temperature | -1.33 [-2.30, -0.39] | -0.30 [-0.46, -0.10] | -2.93 | **0.004** | 0.44 [0.00, 0.66] |
| CV precipitation | -4.43 [-8.14, -0.72] | -0.25 [-0.42, -0.05] | -2.43 | **0.017** | 0.42 [0.00, 0.64] |
| P2T | 0.22 [-0.13, 0.53] | 0.14 [-0.07, 0.33] | 1.33 | 0.188 | 0.53 [0.00, 0.73] |

**Table S5.** Associations between species-specific environmental effects and male body size. Associations of either snout-vent length or body mass as two measures of body size with brumation duration and different environmental variables. All analyses based on df = 114, and confidence intervals are bootstrapped (*N* = 100 simulations). CV = coefficient of variation, P2T = duration of the dry season (in months), with a month defined as dry when its total precipitation is less than two times the mean temperature, and λ = phylogenetic scaling parameter.

| **Predictor** | ***β* [95%CI]** | ***r* [95%CI]** | ***t*** | ***P*** | ***λ* [95%CI]** |
|---|---|---|---|---|---|
| *Snout-vent length:* | | | | | |
| Brumation duration | 0.02 [-0.04, 0.07] | 0.06 [-0.13, 0.23] | 0.59 | 0.557 | 0.93 [0.82, 0.99] |
| Latitude | 0.04 [-0.03, 0.09] | 0.13 [-0.06, 0.30] | 1.36 | 0.177 | 0.93 [0.82, 0.99] |
| Elevation | -0.01 [-0.07, 0.03] | -0.05 [-0.23, 0.13] | -0.53 | 0.597 | 0.94 [0.85, 1.00] |
| Mean temperature | -0.01 [-0.05, 0.05] | -0.04 [-0.21, 0.15] | -0.38 | 0.702 | 0.93 [0.82, 0.99] |
| CV temperature | 0.01 [-0.05, 0.06] | 0.04 [-0.14, 0.22] | 0.42 | 0.676 | 0.93 [0.82, 0.99] |
| P2T | 0.01 [-0.02, 0.06] | 0.05 [-0.13, 0.23] | 0.55 | 0.583 | 0.94 [0.83, 0.99] |
| *Body mass:* | | | | | |
| Brumation duration | -0.03 [-0.23, 0.14] | -0.03 [-0.21, 0.15] | -0.36 | 0.722 | 0.92 [0.80, 0.98] |
| Latitude | 0.10 [-0.12, 0.27] | 0.09 [-0.09, 0.27] | 1.00 | 0.321 | 0.91 [0.77, 0.97] |
| Elevation | -0.16 [-0.32, -0.02] | -0.18 [-0.34, 0.01] | -1.91 | 0.058 | 0.94 [0.85, 1.00] |
| Mean temperature | 0.03 [-0.11, 0.23] | 0.03 [-0.15, 0.21] | 0.37 | 0.710 | 0.92 [0.79, 0.99] |
| CV temperature | -0.03 [-0.22, 0.12] | -0.03 [-0.21, 0.15] | -0.36 | 0.721 | 0.92 [0.79, 0.99] |
| P2T | 0.14 [0.02, 0.29] | 0.16 [-0.02, 0.33] | 1.72 | 0.087 | 0.93 [0.81, 0.98] |

**Table S6.** Associations between tissue size and body size. Log-log association of tissue mass with snout-vent length. To facilitate the interpretation of the allometric slopes (*β*), SVL is cubed ($SVL^3$) so that *β* = 1 equals isometry. All analyses are based on df = 114, and confidence intervals are bootstrapped (*N* = 100 simulations). λ = phylogenetic scaling parameter.

| **Response** | ***β* [95%CI]** | ***r* [95%CI]** | ***t*** | ***P*** | ***λ* [95%CI]** |
|---|---|---|---|---|---|
| Brain | **0.49 [0.44, 0.54]** | 0.88 [0.84, 0.91] | 19.66 | **<0.001** | 0.73 [0.35, 0.87] |
| Body fat | **1.36 [1.28, 1.48]** | 0.94 [0.92, 0.95] | 29.50 | **<0.001** | 0.39 [0.00, 0.64] |
| Digestive tract | 0.95 [0.88, 1.01] | 0.91 [0.88, 0.93] | 23.60 | **<0.001** | 0.00 [0.00, 0.10] |
| Heart | 1.07 [0.98, 1.17] | 0.89 [0.86, 0.92] | 21.27 | **<0.001** | 0.18 [0.00, 0.45] |
| Lungs | 1.06 [0.96, 1.19] | 0.87 [0.83, 0.90] | 18.81 | **<0.001** | 0.23 [0.00, 0.50] |
| Liver | 1.03 [0.96, 1.10] | 0.93 [0.91, 0.95] | 27.12 | **<0.001** | 0.00 [0.00, 0.10] |
| Kidneys | 1.03 [0.97, 1.09] | 0.95 [0.93, 0.96] | 30.90 | **<0.001** | 0.00 [0.00, 0.10] |
| Spleen | 1.10 [0.98, 1.26] | 0.84 [0.79, 0.88] | 16.56 | **<0.001** | 0.38 [0.00, 0.64] |
| Forelimb muscles | **1.23 [1.16, 1.30]** | 0.94 [0.93, 0.96] | 30.62 | **<0.001** | 0.00 [0.00, 0.10] |
| Hindlimb muscles | **1.10 [1.03, 1.18]** | 0.95 [0.93, 0.96] | 31.01 | **<0.001** | 0.49 [0.00, 0.71] |
| Testes | 1.06 [0.94, 1.22] | 0.82 [0.76, 0.86] | 15.27 | **<0.001** | 0.56 [0.00, 0.77] |

**Table S7.** Results of phylogenetically controlled relationships (λ = phylogenetic scaling parameter) between the brumation duration and the relative size of different male tissues in post-brumation breeding (*N* = 116 species) or pre-brumation condition (*N* = 50 species), measured as tissue mass. All associations are controlled for snout-vent length (SVL), and confidence intervals are bootstrapped (*N* = 100 simulations).

| Response | Predictors | *β* [95%CI] | Partial *r* [95%CI] | *t* | *P* | *λ* [95%CI] |
|---|---|---|---|---|---|---|
| *Males in breeding condition (df = 113)* | | | | | | |
| Brain | Brumation | -0.09 [-0.15, -0.04] | -0.31 [-0.46, -0.14] | -3.50 | **<0.001** | 0.35 [0.00, 0.61] |
| | SVL | 0.65 [ 0.59, 0.71] | 0.91 [ 0.87, 0.93] | 22.76 | **<0.001** | |
| Body fat | Brumation | 0.20 [0.04, 0.34] | 0.25 [0.07, 0.40] | 2.72 | **0.008** | 0.42 [0.00, 0.66] |
| | SVL | 1.55 [1.37, 1.73] | 0.87 [0.83, 0.90] | 18.72 | **<0.001** | |
| Digestive tract | Brumation | 0.03 [-0.11, 0.14] | 0.04 [-0.14, 0.22] | 0.45 | 0.656 | 0.00 [0.00, 0.11] |
| | SVL | 1.12 [ 1.01, 1.25] | 0.86 [ 0.81, 0.89] | 17.66 | **<0.001** | |
| Heart | Brumation | -0.07 [-0.22, 0.06] | -0.09 [-0.26, 0.09] | -0.96 | 0.340 | 0.23 [0.00, 0.51] |
| | SVL | 1.35 [ 1.20, 1.50] | 0.87 [ 0.82, 0.90] | 18.49 | **<0.001** | |
| Lungs | Brumation | -0.05 [-0.23, 0.09] | -0.06 [-0.24, 0.12] | -0.69 | 0.490 | 0.21 [0.00, 0.50] |
| | SVL | 1.31 [ 1.15, 1.49] | 0.83 [ 0.77, 0.87] | 15.86 | **<0.001** | |
| Liver | Brumation | 0.05 [-0.09, 0.16] | 0.07 [-0.11, 0.25] | 0.77 | 0.445 | 0.00 [0.00, 0.11] |
| | SVL | 1.23 [ 1.11, 1.36] | 0.87 [ 0.83, 0.90] | 19.12 | **<0.001** | |
| Kidneys | Brumation | -0.02 [-0.14, 0.08] | -0.03 [-0.21, 0.15] | -0.37 | 0.715 | 0.00 [0.00, 0.11] |
| | SVL | 1.25 [ 1.15, 1.36] | 0.90 [ 0.87, 0.92] | 22.15 | **<0.001** | |
| Spleen | Brumation | -0.12 [-0.31, 0.04] | -0.12 [-0.30, 0.06] | -1.32 | 0.189 | 0.23 [0.00, 0.52] |
| | SVL | 1.34 [ 1.15, 1.55] | 0.80 [ 0.74, 0.85] | 14.27 | **<0.001** | |
| Foreleg muscles | Brumation | -0.06 [-0.20, 0.05] | -0.09 [-0.27, 0.09] | -0.98 | 0.329 | 0.00 [0.00, 0.11] |
| | SVL | 1.52 [ 1.40, 1.66] | 0.91 [ 0.88, 0.93] | 23.29 | **<0.001** | |
| Hindleg muscles | Brumation | -0.13 [-0.25, -0.03] | -0.22 [-0.38, -0.04] | -2.37 | **0.019** | 0.22 [0.00, 0.51] |
| | SVL | 1.35 [ 1.23, 1.48] | 0.91 [ 0.88, 0.93] | 22.98 | **<0.001** | |
| Testes | Brumation | 0.30 [0.15, 0.44] | 0.36 [0.19, 0.50] | 4.06 | **<0.001** | 0.77 [0.40, 0.90] |
| | SVL | 1.21 [1.01, 1.41] | 0.78 [0.71, 0.83] | 13.30 | **<0.001** | |
| *Males in pre-brumation condition (df = 47)* | | | | | | |
| Brain | Brumation | -0.15 [-0.21, -0.09] | -0.51 [-0.67, -0.27] | -4.03 | **<0.001** | 0.00 [0.00, 0.56] |
| | SVL | 0.60 [ 0.55, 0.66] | 0.92 [ 0.88, 0.95] | 16.30 | **<0.001** | |
| Body fat | Brumation | 0.33 [0.21, 0.46] | 0.53 [0.29, 0.68] | 4.26 | **<0.001** | 0.55 [0.00, 0.88] |
| | SVL | 1.21 [1.04, 1.40] | 0.89 [0.82, 0.92] | 13.21 | **<0.001** | |
| Digestive tract | Brumation | 0.08 [-0.04, 0.20] | 0.15 [-0.14, 0.40] | 1.01 | 0.320 | 0.64 [0.00, 0.92] |
| | SVL | 1.16 [ 0.99, 1.33] | 0.88 [ 0.81, 0.92] | 12.85 | **<0.001** | |
| Heart | Brumation | -0.02 [-0.15, 0.11] | -0.03 [-0.30, 0.25] | -0.22 | 0.825 | 0.24 [0.00, 0.74] |
| | SVL | 0.89 [ 0.74, 1.07] | 0.83 [ 0.73, 0.88] | 10.11 | **<0.001** | |
| Lungs | Brumation | 0.02 [-0.14, 0.18] | 0.03 [-0.24, 0.31] | 0.24 | 0.812 | 0.55 [0.00, 0.89] |
| | SVL | 1.03 [ 0.81, 1.26] | 0.80 [ 0.68, 0.86] | 9.02 | **<0.001** | |
| Liver | Brumation | 0.11 [-0.02, 0.23] | 0.19 [-0.09, 0.44] | 1.34 | 0.186 | 0.00 [0.00, 0.56] |
| | SVL | 1.16 [ 1.05, 1.30] | 0.90 [ 0.85, 0.93] | 14.56 | **<0.001** | |
| Kidneys | Brumation | -0.06 [-0.17, 0.04] | -0.14 [-0.39, 0.15] | -0.97 | 0.338 | 0.00 [0.00, 0.56] |
| | SVL | 0.99 [ 0.90, 1.11] | 0.91 [ 0.85, 0.94] | 14.85 | **<0.001** | |
| Spleen | Brumation | 0.11 [-0.10, 0.32] | 0.12 [-0.16, 0.38] | 0.83 | 0.410 | 0.00 [0.00, 0.56] |
| | SVL | 1.11 [ 0.93, 1.35] | 0.77 [ 0.64, 0.85] | 8.35 | **<0.001** | |
| Foreleg muscles | Brumation | -0.07 [-0.18, 0.05] | -0.14 [-0.39, 0.15] | -0.94 | 0.354 | 0.00 [0.00, 0.56] |
| | SVL | 1.32 [ 1.22, 1.45] | 0.93 [ 0.90, 0.96] | 18.00 | **<0.001** | |
| Hindleg muscles | Brumation | -0.09 [-0.18, 0.01] | -0.21 [-0.45, 0.07] | -1.49 | 0.142 | 0.57 [0.00, 0.89] |
| | SVL | 1.13 [ 1.01, 1.27] | 0.93 [ 0.88, 0.95] | 16.96 | **<0.001** | |
| Testes | Brumation | 0.02 [-0.14, 0.22] | 0.02 [-0.26, 0.29] | 0.15 | 0.880 | 0.82 [0.15, 0.98] |
| | SVL | 1.00 [ 0.72, 1.24] | 0.73 [ 0.58, 0.82] | 7.34 | **<0.001** | |

**Table S8.** Results of phylogenetically controlled relationships (λ = phylogenetic scaling parameter) of brumation on the relative mass of male tissues (*N* = 116 species). For justification of assignment of brumating and non-brumating species, see Fig. S7. All associations are controlled for snout-vent length (SVL), and confidence intervals are bootstrapped (*N* = 100 simulations).

| Response | Predictors | *β* [95%CI] | Partial *r* [95%CI] | *t* | *P* | *λ* [95%CI] |
|---|---|---|---|---|---|---|
| *Binary classification of brumation (N = 91 brumating, 25 non-brumating species; total df = 113)* | | | | | | |
| Brain | Brumation | -0.12 [-0.25, 0.03] | -0.17 [-0.34, 0.01] | -1.86 | 0.066 | 0.34 [0.00, 0.60] |
| | SVL | 0.64 [ 0.58, 0.71] | 0.90 [ 0.86, 0.92] | 21.56 | **<0.001** | |
| Body fat | Brumation | 0.53 [0.21, 0.93] | 0.26 [0.08, 0.42] | 2.91 | **0.004** | 0.50 [0.00, 0.72] |
| | SVL | 1.53 [1.34, 1.71] | 0.86 [0.82, 0.89] | 18.26 | **<0.001** | |
| Digestive tract | Brumation | 0.01 [-0.30, 0.36] | 0.00 [-0.18, 0.19] | 0.04 | 0.965 | 0.00 [0.00, 0.10] |
| | SVL | 1.13 [ 1.02, 1.27] | 0.86 [ 0.81, 0.89] | 17.64 | **<0.001** | |
| Heart | Brumation | -0.22 [-0.56, 0.16] | -0.12 [-0.30, 0.06] | -1.34 | 0.184 | 0.21 [0.00, 0.49] |
| | SVL | 1.36 [ 1.21, 1.51] | 0.87 [ 0.82, 0.90] | 18.61 | **<0.001** | |
| Lungs | Brumation | -0.07 [-0.47, 0.36] | -0.04 [-0.22, 0.15] | -0.39 | 0.697 | 0.20 [0.00, 0.48] |
| | SVL | 1.31 [ 1.14, 1.48] | 0.83 [ 0.77, 0.87] | 15.71 | **<0.001** | |
| Liver | Brumation | 0.03 [-0.28, 0.39] | 0.02 [-0.16, 0.20] | 0.21 | 0.833 | 0.00 [0.00, 0.10] |
| | SVL | 1.24 [ 1.13, 1.38] | 0.87 [ 0.83, 0.90] | 19.11 | **<0.001** | |
| Kidneys | Brumation | -0.11 [-0.38, 0.20] | -0.08 [-0.25, 0.11] | -0.83 | 0.408 | 0.00 [0.00, 0.10] |
| | SVL | 1.26 [ 1.16, 1.38] | 0.90 [ 0.87, 0.92] | 22.20 | **<0.001** | |
| Spleen | Brumation | -0.39 [-0.83, 0.10] | -0.17 [-0.34, 0.01] | -1.83 | 0.071 | 0.23 [0.00, 0.51] |
| | SVL | 1.36 [ 1.17, 1.55] | 0.81 [ 0.74, 0.85] | 14.47 | **<0.001** | |
| Foreleg muscles | Brumation | -0.07 [-0.39, 0.29] | -0.04 [-0.22, 0.14] | -0.45 | 0.653 | 0.00 [0.00, 0.10] |
| | SVL | 1.51 [ 1.39, 1.66] | 0.91 [ 0.88, 0.93] | 22.87 | **<0.001** | |
| Hindleg muscles | Brumation | -0.15 [-0.44, 0.16] | -0.11 [-0.28, 0.08] | -1.12 | 0.264 | 0.17 [0.00, 0.45] |
| | SVL | 1.34 [ 1.22, 1.44] | 0.90 [ 0.87, 0.93] | 22.41 | **<0.001** | |
| Testes | Brumation | 0.49 [ 0.13, 0.89] | 0.23 [ 0.05, 0.39] | 2.49 | **0.014** | 0.75 [0.36, 0.90] |
| | SVL | 1.23 [ 1.02, 1.43] | 0.77 [ 0.70, 0.82] | 12.91 | **<0.001** | |
| *Binary classification of brumation with a 4°C buffer (N = 47 brumating, 69 non-brumating species; total df = 113)* | | | | | | |
| Brain | Brumation | -0.21 [-0.31, -0.14] | -0.38 [-0.51, -0.21] | -4.33 | **<0.001** | 0.35 [0.00, 0.62] |
| | SVL | 0.64 [ 0.59, 0.71] | 0.91 [ 0.88, 0.93] | 23.47 | **<0.001** | |
| Body fat | Brumation | 0.29 [ 0.00, 0.52] | 0.19 [ 0.01, 0.36] | 2.07 | **0.041** | 0.46 [0.00, 0.72] |
| | SVL | 1.57 [ 1.39, 1.77] | 0.87 [ 0.83, 0.90] | 18.88 | **<0.001** | |
| Digestive tract | Brumation | 0.07 [-0.17, 0.23] | 0.06 [-0.13, 0.23] | 0.60 | 0.553 | 0.00 [0.00, 0.11] |
| | SVL | 1.12 [ 1.01, 1.24] | 0.87 [ 0.82, 0.90] | 18.43 | **<0.001** | |
| Heart | Brumation | -0.25 [-0.52, -0.07] | -0.18 [-0.35, 0.00] | -1.97 | 0.051 | 0.23 [0.00, 0.51] |
| | SVL | 1.35 [ 1.21, 1.52] | 0.88 [ 0.83, 0.90] | 19.23 | **<0.001** | |
| Lungs | Brumation | -0.31 [-0.61, -0.10] | -0.20 [-0.36, -0.01] | -2.13 | **0.035** | 0.21 [0.00, 0.50] |
| | SVL | 1.32 [ 1.17, 1.51] | 0.84 [ 0.79, 0.88] | 16.68 | **<0.001** | |
| Liver | Brumation | -0.09 [-0.34, 0.06] | -0.07 [-0.25, 0.11] | -0.76 | 0.45 | 0.00 [0.00, 0.11] |
| | SVL | 1.25 [ 1.14, 1.38] | 0.89 [ 0.85, 0.91] | 20.31 | **<0.001** | |
| Kidneys | Brumation | -0.06 [-0.27, 0.08] | -0.05 [-0.23, 0.13] | -0.55 | 0.586 | 0.00 [0.00, 0.11] |
| | SVL | 1.24 [ 1.15, 1.35] | 0.91 [ 0.88, 0.93] | 23.04 | **<0.001** | |
| Spleen | Brumation | -0.19 [-0.54, 0.05] | -0.10 [-0.28, 0.08] | -1.12 | 0.265 | 0.23 [0.00, 0.51] |
| | SVL | 1.33 [ 1.14, 1.54] | 0.80 [ 0.74, 0.85] | 14.42 | **<0.001** | |
| Foreleg muscles | Brumation | -0.26 [-0.50, -0.10] | -0.19 [-0.35, -0.01] | -2.04 | **0.043** | 0.00 [0.00, 0.11] |
| | SVL | 1.52 [ 1.41, 1.65] | 0.92 [ 0.89, 0.94] | 24.62 | **<0.001** | |
| Hindleg muscles | Brumation | -0.23 [-0.45, -0.08] | -0.20 [-0.36, -0.02] | -2.16 | **0.033** | 0.24 [0.00, 0.52] |
| | SVL | 1.33 [ 1.22, 1.47] | 0.91 [ 0.88, 0.93] | 23.04 | **<0.001** | |
| Testes | Brumation | 0.38 [ 0.09, 0.60] | 0.24 [ 0.06, 0.40] | 2.63 | **0.01** | 0.74 [0.34, 0.89] |
| | SVL | 1.25 [ 1.03, 1.44] | 0.78 [ 0.71, 0.83] | 13.31 | **<0.001** | |

**Table S9.** Effects of the brumation duration on relative tissue size in males, excluding those 25 species that are unlikely to overwinter (≤27 days). Results of phylogenetically controlled relationships (λ = phylogenetic scaling parameter) between the brumation duration and the relative mass of different male tissues in breeding condition (df = 88). All associations are controlled for snout-vent length (SVL), and confidence intervals are bootstrapped (*N* = 100 simulations).

| **Response** | **Predictors** | ***β* [95%CI]** | **Partial *r* [95%CI]** | ***t*** | ***P*** | ***λ* [95%CI]** |
|---|---|---|---|---|---|---|
| Brain | Brumation | -0.07 [-0.13, -0.02] | -0.27 [-0.44, -0.07] | -2.62 | **0.010** | 0.49 [0.00, 0.72] |
| | SVL | 0.56 [ 0.50, 0.62] | 0.88 [ 0.84, 0.91] | 17.52 | **<0.001** | |
| Body fat | Brumation | 0.07 [-0.08, 0.22] | 0.10 [-0.11, 0.29] | 0.92 | 0.36 | 0.38 [0.00, 0.61] |
| | SVL | 1.43 [ 1.26, 1.60] | 0.88 [ 0.83, 0.91] | 17.08 | **<0.001** | |
| Digestive tract | Brumation | 0.04 [-0.09, 0.20] | 0.06 [-0.15, 0.26] | 0.55 | 0.587 | 0.00 [0.00, 0.11] |
| | SVL | 0.99 [ 0.84, 1.13] | 0.83 [ 0.77, 0.87] | 14.03 | **<0.001** | |
| Heart | Brumation | -0.07 [-0.20, 0.09] | -0.10 [-0.29, 0.11] | -0.93 | 0.355 | 0.13 [0.00, 0.33] |
| | SVL | 1.28 [ 1.14, 1.44] | 0.88 [ 0.83, 0.91] | 17.14 | **<0.001** | |
| Lungs | Brumation | -0.11 [-0.27, 0.06] | -0.15 [-0.34, 0.06] | -1.40 | 0.165 | 0.23 [0.00, 0.46] |
| | SVL | 1.24 [ 1.08, 1.42] | 0.84 [ 0.78, 0.88] | 14.75 | **<0.001** | |
| Liver | Brumation | 0.00 [-0.14, 0.16] | -0.01 [-0.21, 0.20] | -0.06 | 0.949 | 0.05 [0.00, 0.19] |
| | SVL | 1.13 [ 0.99, 1.29] | 0.85 [ 0.80, 0.89] | 15.38 | **<0.001** | |
| Kidneys | Brumation | 0.03 [-0.08, 0.16] | 0.05 [-0.16, 0.25] | 0.48 | 0.635 | 0.00 [0.00, 0.11] |
| | SVL | 1.15 [ 1.03, 1.27] | 0.90 [ 0.87, 0.93] | 19.88 | **<0.001** | |
| Spleen | Brumation | -0.06 [-0.23, 0.13] | -0.07 [-0.26, 0.14] | -0.61 | 0.541 | 0.29 [0.00, 0.52] |
| | SVL | 1.32 [ 1.12, 1.53] | 0.82 [ 0.75, 0.86] | 13.32 | **<0.001** | |
| Foreleg muscles | Brumation | -0.10 [-0.23, 0.06] | -0.16 [-0.35, 0.05] | -1.49 | 0.139 | 0.00 [0.00, 0.11] |
| | SVL | 1.41 [ 1.26, 1.55] | 0.91 [ 0.87, 0.93] | 20.34 | **<0.001** | |
| Hindleg muscles | Brumation | -0.16 [-0.27, -0.04] | -0.30 [-0.47, -0.10] | -2.93 | **0.004** | 0.15 [0.00, 0.35] |
| | SVL | 1.27 [ 1.16, 1.40] | 0.92 [ 0.89, 0.94] | 21.76 | **<0.001** | |
| Testes | Brumation | 0.19 [ 0.02, 0.35] | 0.25 [0.04, 0.42] | 2.38 | **0.019** | 0.55 [0.03, 0.76] |
| | SVL | 1.21 [ 1.01, 1.39] | 0.80 [0.72, 0.85] | 12.56 | **<0.001** | |

**Table S10.** Effects of brumation on relative tissue size in males, using different temperature thresholds. Phylogenetically controlled relationships (λ = phylogenetic scaling parameter) between the brumation period and the relative male tissue mass in breeding condition (*N* = 116 species), with a 2 or 4°C buffer relative to Table S7. All associations are controlled for snout-vent length, confidence intervals are bootstrapped (*N* = 100 simulations).

| Response | Predictors | *β* [95%CI] | Partial *r* [95%CI] | *t* | *P* | *λ* [95%CI] |
|---|---|---|---|---|---|---|
| *Brumation with 2°C temperature buffer (df = 113)* | | | | | | |
| Brain | Brumation | -0.08 [-0.14, -0.04] | -0.30 [-0.45, -0.12] | -3.29 | **0.001** | 0.35 [0.00, 0.61] |
| | SVL | 0.65 [ 0.59, 0.71] | 0.91 [ 0.87, 0.93] | 22.65 | **<0.001** | |
| Body fat | Brumation | 0.17 [ 0.02, 0.29] | 0.21 [ 0.03, 0.37] | 2.29 | **0.024** | 0.44 [0.00, 0.69] |
| | SVL | 1.56 [ 1.38, 1.75] | 0.87 [ 0.83, 0.90] | 18.79 | **<0.001** | |
| Digestive tract | Brumation | 0.03 [-0.09, 0.12] | 0.04 [-0.14, 0.22] | 0.43 | 0.667 | 0.00 [0.00, 0.11] |
| | SVL | 1.12 [ 1.01, 1.25] | 0.86 [ 0.81, 0.89] | 17.91 | **<0.001** | |
| Heart | Brumation | -0.10 [-0.24, 0.01] | -0.14 [-0.31, 0.05] | -1.45 | 0.15 | 0.23 [0.00, 0.51] |
| | SVL | 1.35 [ 1.21, 1.51] | 0.87 [ 0.83, 0.90] | 18.84 | **<0.001** | |
| Lungs | Brumation | -0.09 [-0.25, 0.03] | -0.11 [-0.29, 0.07] | -1.22 | 0.226 | 0.23 [0.00, 0.51] |
| | SVL | 1.32 [ 1.16, 1.50] | 0.83 [ 0.78, 0.87] | 16.13 | **<0.001** | |
| Liver | Brumation | 0.01 [-0.11, 0.11] | 0.02 [-0.17, 0.20] | 0.17 | 0.867 | 0.00 [0.00, 0.11] |
| | SVL | 1.24 [ 1.13, 1.37] | 0.88 [ 0.84, 0.91] | 19.53 | **<0.001** | |
| Kidneys | Brumation | -0.01 [-0.11, 0.08] | -0.01 [-0.19, 0.17] | -0.10 | 0.924 | 0.00 [0.00, 0.11] |
| | SVL | 1.24 [ 1.14, 1.36] | 0.90 [ 0.87, 0.92] | 22.33 | **<0.001** | |
| Spleen | Brumation | -0.12 [-0.30, 0.02] | -0.13 [-0.30, 0.05] | -1.39 | 0.167 | 0.24 [0.00, 0.52] |
| | SVL | 1.34 [ 1.16, 1.55] | 0.80 [ 0.74, 0.85] | 14.38 | **<0.001** | |
| Foreleg muscles | Brumation | -0.09 [-0.21, 0.01] | -0.13 [-0.30, 0.05] | -1.43 | 0.156 | 0.00 [0.00, 0.11] |
| | SVL | 1.53 [ 1.42, 1.66] | 0.91 [ 0.88, 0.93] | 23.80 | **<0.001** | |
| Hindleg muscles | Brumation | -0.16 [-0.28, -0.08] | -0.28 [-0.43, -0.10] | -3.07 | **0.003** | 0.25 [0.00, 0.54] |
| | SVL | 1.35 [ 1.24, 1.48] | 0.91 [ 0.88, 0.93] | 23.46 | **<0.001** | |
| Testes | Brumation | 0.25 [ 0.12, 0.39] | 0.31 [ 0.13, 0.46] | 3.46 | **<0.001** | 0.75 [0.37, 0.89] |
| | SVL | 1.23 [ 1.02, 1.42] | 0.78 [ 0.71, 0.83] | 13.31 | **<0.001** | |
| *Brumation with 4°C temperature buffer (df = 113)* | | | | | | |
| Brain | Brumation | -0.09 [-0.14, -0.05] | -0.33 [-0.47, -0.16] | -3.72 | **<0.001** | 0.34 [0.00, 0.61] |
| | SVL | 0.65 [ 0.59, 0.71] | 0.91 [ 0.88, 0.93] | 23.01 | **<0.001** | |
| Body fat | Brumation | 0.14 [ 0.00, 0.26] | 0.17 [-0.01, 0.34] | 1.86 | 0.065 | 0.44 [0.00, 0.69] |
| | SVL | 1.56 [ 1.38, 1.76] | 0.87 [ 0.82, 0.90] | 18.63 | **<0.001** | |
| Digestive tract | Brumation | 0.03 [-0.08, 0.12] | 0.05 [-0.14, 0.22] | 0.48 | 0.63 | 0.00 [0.00, 0.11] |
| | SVL | 1.12 [ 1.02, 1.25] | 0.86 [ 0.81, 0.89] | 17.98 | **<0.001** | |
| Heart | Brumation | -0.07 [-0.19, 0.04] | -0.10 [-0.27, 0.09] | -1.05 | 0.295 | 0.23 [0.00, 0.52] |
| | SVL | 1.35 [ 1.21, 1.50] | 0.87 [ 0.83, 0.90] | 18.67 | **<0.001** | |
| Lungs | Brumation | -0.12 [-0.26, 0.00] | -0.15 [-0.32, 0.03] | -1.66 | 0.099 | 0.24 [0.00, 0.53] |
| | SVL | 1.33 [ 1.17, 1.51] | 0.84 [ 0.78, 0.87] | 16.25 | **<0.001** | |
| Liver | Brumation | 0.01 [-0.11, 0.10] | 0.01 [-0.17, 0.19] | 0.10 | 0.92 | 0.00 [0.00, 0.11] |
| | SVL | 1.24 [ 1.14, 1.37] | 0.88 [ 0.84, 0.91] | 19.63 | **<0.001** | |
| Kidneys | Brumation | 0.00 [-0.10, 0.08] | -0.01 [-0.19, 0.18] | -0.06 | 0.956 | 0.00 [0.00, 0.11] |
| | SVL | 1.24 [ 1.15, 1.35] | 0.90 [ 0.87, 0.93] | 22.41 | <0.001 | |
| Spleen | Brumation | -0.11 [-0.26, 0.03] | -0.12 [-0.29, 0.07] | -1.25 | 0.213 | 0.23 [0.00, 0.52] |
| | SVL | 1.34 [ 1.16, 1.54] | 0.80 [ 0.74, 0.85] | 14.34 | **<0.001** | |
| Foreleg muscles | Brumation | -0.10 [-0.21, -0.01] | -0.15 [-0.32, 0.04] | -1.56 | 0.121 | 0.00 [0.00, 0.11] |
| | SVL | 1.53 [ 1.42, 1.66] | 0.91 [ 0.89, 0.93] | 23.95 | **<0.001** | |
| Hindleg muscles | Brumation | -0.15 [-0.25, -0.07] | -0.26 [-0.42, -0.09] | -2.91 | **0.004** | 0.27 [0.00, 0.55] |
| | SVL | 1.35 [ 1.23, 1.48] | 0.91 [ 0.88, 0.93] | 23.24 | **<0.001** | |
| Testes | Brumation | 0.22 [ 0.08, 0.34] | 0.28 [ 0.10, 0.43] | 3.10 | **0.002** | 0.76 [0.39, 0.90] |
| | SVL | 1.22 [ 1.01, 1.42] | 0.78 [ 0.70, 0.83] | 13.05 | **<0.001** | |

**Table S11.** Species-specific environmental effects on the duration of the breeding season. CV = coefficient of variation, P2T = duration of the dry season (in months), with a month defined as dry when its total precipitation is less than two times the mean temperature. All analyses based on df = 41. The regression slope *β* is presented with its bootstrapped confidence interval, λ is the phylogenetic scaling parameter.

| Predictor | *β* [95%CI] | *r* [95%CI] | *t* | *P* | *λ* [95%CI] |
|---|---|---|---|---|---|
| Brumation | -8.12 [-11.70, -5.01] | -0.57 [-0.72, -0.33] | -4.47 | **<0.001** | 0.00 [0.00, 0.38] |
| Latitude | -4.86 [-8.95, -1.50] | -0.34 [-0.56, -0.05] | -2.33 | **0.025** | 0.00 [0.00, 0.39] |
| Longitude | 3.43 [-0.94, 8.44] | 0.24 [-0.06, 0.49] | 1.59 | 0.119 | 0.00 [0.00, 0.37] |
| Elevation | -5.56 [-9.84, -1.61] | -0.39 [-0.60, -0.10] | -2.73 | **0.009** | 0.00 [0.00, 0.38] |
| Mean temperature | 5.65 [1.78, 9.93] | 0.40 [0.11, 0.60] | 2.78 | **0.008** | 0.00 [0.00, 0.31] |
| Mean precipitation | 3.62 [-0.75, 7.62] | 0.26 [-0.05, 0.50] | 1.69 | 0.098 | 0.00 [0.00, 0.40] |
| CV temperature | -6.06 [-10.14, -2.06] | -0.43 [-0.62, -0.15] | -3.03 | **0.004** | 0.00 [0.00, 0.42] |
| CV precipitation | -1.44 [-5.70, 2.87] | -0.10 [-0.38, 0.20] | -0.65 | 0.518 | 0.00 [0.00, 0.42] |
| P2T | 2.73 [-1.28, 7.59] | 0.19 [-0.11, 0.45] | 1.25 | 0.217 | 0.00 [0.00, 0.40] |

**Table S12.** Species-specific environmental effects on the duration of the breeding season when accounting for brumation duration. CV = coefficient of variation, P2T = duration of the dry season (in months), with a month defined as dry when its total precipitation is less than two times the mean temperature. All analyses based on df = 40. The regression slope *β* is presented with its bootstrapped confidence interval, λ is the phylogenetic scaling parameter.

| **Predictors** | ***β* [95%CI]** | ***r* [95%CI]** | ***t*** | ***P*** | ***λ* [95%CI]** |
|---|---|---|---|---|---|
| Latitude | 2.01 [ -2.72, 6.60] | 0.12 [-0.19, 0.40] | 0.77 | 0.448 | 0.00 [0.00, 0.40] |
| Brumation | -9.56 [-14.76, -4.87] | -0.50 [-0.67, -0.23] | -3.64 | **<0.001** | |
| Longitude | 1.35 [ -2.38, 4.50] | 0.11 [-0.20, 0.39] | 0.71 | 0.482 | 0.00 [0.00, 0.34] |
| Brumation | -7.76 [-10.87, -4.37] | -0.54 [-0.70, -0.29] | -4.09 | **<0.001** | |
| Elevation | -2.12 [ -5.76, 1.96] | -0.16 [-0.43, 0.15] | -1.02 | 0.313 | 0.00 [0.00, 0.57] |
| Brumation | -7.09 [-12.59, -3.16] | -0.48 [-0.66, -0.20] | -3.41 | **0.001** | |
| Temperature | 2.45 [ -1.38, 5.77] | 0.19 [-0.12, 0.45] | 1.20 | 0.235 | 0.00 [0.00, 0.46] |
| Brumation | -7.00 [-11.08, -3.04] | -0.48 [-0.66, -0.21] | -3.44 | **0.001** | |
| Precipitation | 0.23 [ -3.31, 3.77] | 0.02 [-0.28, 0.31] | 0.12 | 0.909 | 0.00 [0.00, 0.27] |
| Brumation | -8.02 [-11.26, -4.34] | -0.53 [-0.69, -0.27] | -3.95 | **<0.001** | |
| CV temperature | -2.41 [ -6.91, 1.41] | -0.18 [-0.44, 0.13] | -1.13 | 0.267 | 0.00 [0.00, 0.46] |
| Brumation | -6.83 [-10.85, -2.64] | -0.45 [-0.64, -0.17] | -3.19 | **0.003** | |
| CV precipitation | 0.64 [ -2.76, 3.95] | 0.05 [-0.25, 0.34] | 0.34 | 0.739 | 0.00 [0.00, 0.49] |
| Brumation | -8.28 [-11.95, -4.51] | -0.57 [-0.72, -0.32] | -4.36 | **<0.001** | |
| P2T | 1.20 [ -2.12, 4.46] | 0.10 [-0.21, 0.38] | 0.64 | 0.524 | 0.00 [0.00, 0.44] |
| Brumation | -7.89 [-10.79, -3.66] | -0.56 [-0.71, -0.31] | -4.23 | **<0.001** | |

**Table S13.** Comparison of candidate models in a phylogenetic path analysis linking environmental variation to brumation duration and the breeding patterns (breeding season and aggregations). All candidate models are ranked according to their *C*-statistic Information Criterion (CICc). Those with ΔCICc < 2 are highlighted in bold and were used to calculate the average model, which is depicted in Fig. S4 along with all hypotheses.

| Model | *k* | *q* | *C* | *P* | CICc | ΔCICc | $w_i$ |
|---|---|---|---|---|---|---|---|
| ***m5*** | **3** | **7** | **3.32** | **0.77** | **20.52** | **0.00** | **1.00** |
| ***m4*** | **2** | **8** | **1.72** | **0.79** | **21.95** | **1.43** | **0.49** |
| *m1* | 1 | 9 | 0.29 | 0.87 | 23.74 | 3.22 | 0.20 |
| *m3* | 2 | 8 | 15.06 | 0.00 | 35.29 | 14.77 | 0.00 |
| *m6* | 3 | 7 | 22.03 | 0.00 | 39.23 | 18.71 | 0.00 |
| *m2* | 3 | 7 | 35.60 | 0.00 | 52.80 | 32.28 | 0.00 |
| *m8* | 3 | 7 | 60.09 | 0.00 | 77.29 | 56.77 | 0.00 |
| *m7* | 2 | 8 | 83.01 | 0.00 | 103.25 | 82.73 | 0.00 |

*k* = number of independence claims; *q* = number of parameters; *C* = Fisher's *C* statistics; CICc = *C*-statistic Information Criterion; ΔCICc = difference in CICc from the best-fitting model; $w_i$ = CICc weight.

**Table S14.** Correlation matrix between the compositional data of the four focal traits and the rest of the body. The upper triangle lists the correlation coefficients with 95% confidence intervals (controlling for phylogeny), the lower triangle the corresponding *P*-values before (upper) and after (lower) correction for multiple testing based on the false discovery rate. Statistically significant correlations are highlighted in bold.

| | Brain | Body fat | Testes | Hindlimb muscles | Rest |
|---|---|---|---|---|---|
| Brain | | **-0.79**<br>**[-0.83, -0.71]** | **-0.36**<br>**[-0.50, -0.19]** | -0.08<br>[-0.26, 0.10] | **0.30**<br>**[0.12, 0.44]** |
| Body fat | **<0.001**<br>**<0.001** | | -0.06<br>[-0.23, 0.12] | -0.03<br>[-0.20, 0.15] | -0.17<br>[-0.34, 0.01] |
| Testes | **<0.001**<br>**<0.001** | 0.531<br>0.590 | | **-0.54**<br>**[-0.64, -0.40]** | **-0.57**<br>**[-0.67, -0.44]** |
| Hindlimb muscles | 0.370<br>0.527 | 0.776<br>0.776 | **<0.001**<br>**<0.001** | | 0.07<br>[-0.11, 0.24] |
| Rest | **0.001**<br>**0.002** | 0.060<br>0.101 | **<0.001**<br>**<0.001** | 0.466<br>0.582 | |

**Table S15.** Results of the phylogenetic principal component analysis. Shown are the eigenvalues and percent of the total variance explained by the four principal components, as well as the loadings of the original variables. Loadings >0.25 are highlighted in bold. Phylogenetic scaling parameter $\lambda$ = 0.75.

| | **PC1** | **PC2** | **PC3** | **PC4** |
|---|---|---|---|---|
| Eigenvalues | 3.38 | 0.32 | 0.21 | 0.09 |
| % variance | 84.4 | 8.0 | 5.2 | 2.3 |
| *Loadings* | | | | |
| Brain | **-0.91** | **0.31** | **0.26** | -0.12 |
| Body fat | **-0.93** | -0.07 | **-0.31** | -0.16 |
| Testes | **-0.88** | **-0.44** | 0.18 | 0.04 |
| Hindlimb muscles | **-0.95** | 0.18 | -0.11 | 0.23 |

**Table S16.** Phylogenetic path analysis linking brumation to relative trait investments. All candidate models are ranked according to their CICc. Those with ΔCICc < 2 are highlighted in bold and were used to calculate the average model (see Fig. 4 in main text). For candidate models and averaged path coefficients see Figs. S6 and S7.

| Model | *k* | *q* | *C* | *P* | CICc | ΔCICc | $w_i$ |
|---|---|---|---|---|---|---|---|
| ***m11*** | **4** | **17** | **8.28** | **0.41** | **48.50** | **0.00** | 0.25 |
| ***m25*** | **1** | **20** | **0.99** | **0.61** | **49.80** | **1.31** | 0.13 |
| ***m23*** | **1** | **20** | **1.28** | **0.53** | **50.10** | **1.59** | 0.11 |
| ***m24*** | **1** | **20** | **1.28** | **0.53** | **50.10** | **1.59** | 0.11 |
| ***m3*** | **3** | **18** | **7.44** | **0.28** | **50.50** | **1.96** | 0.09 |
| *m12* | 3 | 18 | 7.53 | 0.28 | 50.60 | 2.05 | 0.09 |
| *m9* | 5 | 16 | 13.48 | 0.20 | 51.00 | 2.45 | 0.07 |
| *m21* | 2 | 19 | 6.52 | 0.16 | 52.40 | 3.91 | 0.03 |
| *m4* | 2 | 19 | 6.68 | 0.15 | 52.60 | 4.08 | 0.03 |
| *m1* | 4 | 17 | 12.64 | 0.13 | 52.90 | 4.35 | 0.03 |
| *m10* | 4 | 17 | 12.72 | 0.12 | 53.00 | 4.44 | 0.03 |
| *m22* | 2 | 19 | 7.54 | 0.11 | 53.50 | 4.93 | 0.02 |
| *m2* | 3 | 18 | 11.88 | 0.06 | 54.90 | 6.41 | 0.01 |
| *m26* | 3 | 18 | 14.45 | 0.03 | 57.50 | 8.98 | 0.00 |
| *m7* | 4 | 17 | 23.32 | 0.00 | 63.60 | 15.04 | 0.00 |
| *m19* | 4 | 17 | 23.95 | 0.00 | 64.20 | 15.66 | 0.00 |
| *m8* | 3 | 18 | 22.57 | 0.00 | 65.60 | 17.09 | 0.00 |
| *m5* | 5 | 16 | 28.52 | 0.00 | 66.00 | 17.49 | 0.00 |
| *m14* | 3 | 18 | 23.59 | 0.00 | 66.60 | 18.11 | 0.00 |
| *m13* | 4 | 17 | 26.53 | 0.00 | 66.80 | 18.25 | 0.00 |
| *m20* | 3 | 18 | 24.03 | 0.00 | 67.10 | 18.56 | 0.00 |
| *m6* | 4 | 17 | 27.76 | 0.00 | 68.00 | 19.48 | 0.00 |
| *m16* | 5 | 16 | 36.26 | 0.00 | 73.80 | 25.23 | 0.00 |
| *m27* | 5 | 16 | 37.41 | 0.00 | 74.90 | 26.38 | 0.00 |
| *m15* | 3 | 18 | 34.57 | 0.00 | 77.60 | 29.10 | 0.00 |
| *m17* | 4 | 17 | 37.68 | 0.00 | 77.90 | 29.39 | 0.00 |
| *m28* | 6 | 15 | 50.57 | 0.00 | 85.40 | 36.85 | 0.00 |
| *m18* | 4 | 17 | 59.08 | 0.00 | 99.30 | 50.80 | 0.00 |

*k* = number of independence claims; *q* = number of parameters; *C* = Fisher's *C* statistics; CICc = *C*-statistic Information Criterion; ΔCICc = difference in CICc from the best-fitting model; $w_i$ = CICc weight.

**Table S17.** Effect of the predominant habitat of each species on the relative amount of body fat in the context of the brain−fat trade-off hypothesis. Output of PGLS models examining the effect of the predominant species-specific habitat (and so mode of locomotion) on the relative amount of body fat, accounting for brain mass and brumation duration as other sources of variation. Three different models were conducted: (1) using raw species means with log body fat as the response variable and controlling for body size by including log snout-vent length as a covariate; (2) using the compositional data (centred log ratio of both brain mass as the response and brain mass); and (3) using PC3 of the principal component analysis, which was primarily loaded by brain mass (+0.26) and body fat (−0.31; Fig. 3). Statistically significant effects are highlighted in bold, and the different habitat types are compared against aquatic species as the reference in the intercept.

| **Predictors** | ***β* [95%CI]** | ***r* [95%CI]** | ***t*** | ***P*** |
|---|---|---|---|---|
| *Raw species means (df = 109, λ = 0.17 [0.00, 0.51])* | | | | |
| (Intercept) | -12.97 [-17.49, -8.41] | -0.44 [-0.57, -0.28] | -5.12 | **<0.001** |
| Log Brain mass | 0.50 [0.12, 0.94] | 0.18 [0.00, 0.35] | 1.94 | 0.055 |
| Habitat [arboreal] | -0.97 [-1.49, -0.38] | -0.3 [-0.45, -0.12] | -3.31 | **0.001** |
| Habitat [semiaquatic] | -0.02 [-0.31, 0.35] | -0.01 [-0.19, 0.18] | -0.09 | 0.925 |
| Habitat [terrestrial] | -0.3 [-0.83, 0.35] | -0.11 [-0.28, 0.08] | -1.12 | 0.264 |
| Brumation | 0.22 [0.09, 0.37] | 0.27 [0.09, 0.42] | 2.90 | **0.005** |
| Log SVL | 2.94 [2.13, 3.76] | 0.52 [0.37, 0.63] | 6.34 | **<0.001** |
| *Compositional data (df = 110, λ = 0.01 [0.00, 0.03])* | | | | |
| (Intercept) | -1.94 [-2.14, -1.69] | -0.86 [-0.89, -0.82] | -17.96 | **<0.001** |
| Brain mass | -0.61 [-0.74, -0.52] | -0.68 [-0.76, -0.58] | -9.81 | **<0.001** |
| Habitat [arboreal] | -0.28 [-0.51, -0.06] | -0.22 [-0.38, -0.04] | -2.39 | **0.018** |
| Habitat [semiaquatic] | -0.10 [-0.31, 0.06] | -0.10 [-0.28, 0.09] | -1.07 | 0.286 |
| Habitat [terrestrial] | 0.01 [-0.19, 0.20] | 0.01 [-0.17, 0.19] | 0.12 | 0.901 |
| Brumation | 0.08 [0.01, 0.17] | 0.19 [0.00, 0.35] | 1.98 | **0.050** |
| *PC3 (df = 111, λ = 0.20 [0.00, 0.59])* | | | | |
| (Intercept) | -0.03 [-0.26, 0.16] | -0.06 [-0.24, 0.12] | -0.27 | 0.785 |
| Habitat [arboreal] | 0.31 [0.12, 0.52] | 0.26 [0.08, 0.42] | 2.83 | **0.006** |
| Habitat [semiaquatic] | -0.01 [-0.15, 0.13] | -0.01 [-0.19, 0.17] | -0.10 | 0.920 |
| Habitat [terrestrial] | 0.10 [-0.06, 0.30] | 0.09 [-0.09, 0.27] | 0.99 | 0.326 |
| Brumation | -0.02 [-0.08, 0.02] | -0.08 [-0.26, 0.10] | -0.88 | 0.381 |
